# In-silico analysis of cyanobacteriochrome architectures and spectral diversity

**DOI:** 10.1101/2022.09.22.509050

**Authors:** Aleksandar Mihnev, Anna Amtmann

**Affiliations:** School of Molecular Biosciences, University of Glasgow, Glasgow G12 8QQ, UK

## Abstract

The cyanobacteriochrome GAF domains represent a trove of spectral diversity. These proteins are endemic to cyanobacteria and sense the color and power of light. Multiple mechanisms are used to tune the natural absorbance spectrum of the bound bilin chromophore. In practice, these are difficult to identify from the predicted amino acid sequence. Their individual presence rarely yields a consistent and predictable outcome. The absorbance characteristics of the GAF domain are a complex function of many such tuning mechanisms. This implies that a more combinatoric approach to characterizing the diversity of GAF domains would better to predict spectral tunes. We reviewed the literature and constructed a dataset of predicted/confirmed cyanobacteriochrome GAF domains. This dataset was subjected to multiple sequence alignments and 18 GAF domain families were defined. The amino acid sequence similarity correlated well with known spectral characteristics but there were exceptions. A second approach to predict chromotype involved using Principal Component Analysis to characterize the whole domain architectures of cyanobacteriochrome. This approach identified 7 conserved domain architectures, with some variations. These also offered a correlation to the spectral tune of the GAF domains therein, in addition to the 18 GAF families. The three-dimensional structures of 98 spectrally characterized GAF domains were predicted using Phyre2. Subsequent grouping based on distance maps offered an insight into how the general spectral position of the domain is set. Finer tuning is likely to be achieved by means of six key residues within the binding pocket. Taken together, these insights allowed us to carry out a Multiple Correlation Analysis serving as a mathematical summary of the diversity of cyanobacteriochrome GAF domains. This summary or “cyanobacteriochrome atlas” can be used to make spectral predictions on uncharacterized GAF domains.

## INTRODUCTION

Cyanobacteriochromes provide important information pertaining to the light environment. These photoreceptor proteins embody a vast spectral diversity (Fushimi and Narikawa, 2019). Despite of this, cyanobacteriochromes (CBCRs) are currently only found in cyanobacteria where they serve to regulate and optimize important light-based processes (Rockwell et al., 2012).

The minimum structural unit of a CBCR is a specific subset of *cGMP-phosphodiesterase/adenlylate cyclase/FhlA* (GAF) domain (Rockwell et al., 2015a). The CBCR GAF domain has evolved to bind a phycocyanobilin (PCB) chromophore (Burgie and Vierstra, 2014; Ikeuchi and Ishizuka, 2008; Rockwell et al., 2011, 2013) which is attuned to the myriad of absorbance spectra (Chen et al., 2012a; Fushimi et al., 2020a; Hirose et al., 2013; Ishizuka et al., 2011; Rockwell et al., 2011, 2013; Xu et al., 2020).

A further level of spectral complexity is added by the fact that PCB can become isomerized upon light excitation, yielding a second absorbance spectrum for the same GAF domain. These two PCB isomers within the protein are referred to as the 15Z and 15E states (Chen et al., 2012a; Rockwell et al., 2013; van Thor et al., 2006).

In many cases the spectral properties of the bilin in solution are different than those of the bilin bound within a CBCR GAF domain. This is due to a number of spectral tuning tricks employed by the protein (Chen et al., 2012b; Fushimi et al., 2020b; Hirose et al., 2013; Ishizuka et al., 2011; Rockwell et al., 2011, 2013; Xu et al., 2020). For these reasons the GAF-PCB holoprotein is often described with P states instead. These reflect the spectral colors most strongly absorbed by the GAF domain in its two states. We define the term “chromotype” to denote the specific combination of spectral colours sensed by a GAF domain, relative to the 15Z/15E chromophore state. For example, a GAF domain with a Pr (15Z) and Pg (15E) configuration will have a Red/Green (R/G) chromotype. Similarly, a Pb (15Z) and Po (15E) GAF domain will have a Blue/Orange (B/O) chromotype.

CBCRs are often more complex than a single GAF domain. The status quo is of CBCRs is usually multi-domain proteins, combination of one or more GAF and other sensory or effector domains. CBCR protein signaling is not as well understood as that in phytochrome or phototropins (Bezy and Kehoe, 2010; Bhaya, 2016; Hirose et al., 2010, 2013; Kehoe and Gutu, 2006). Moreover, only a handful of structural studies on CBCR GAF domains have been done to date.

Our lab is interested in providing a systematic review of the CBCR GAF domain properties available from the literature. This is in the form of a computational summary of the nuances between various CBCR GAF domains. In doing this, we aim to learn from others’ work and provide several directions to fully interrogating the amino acid sequence of uncharacterized GAF domains in the context of predicting chromotype. Chromotype predictions made from genomic data would provide targeted studies of light-induced phenotypes, the means to manipulate cyanobacterial metabolism with light, as well as identifying interesting CBCRs for further studies.

## RESULTS

### Dataset construction

Currently, there are more than 344 thousand putative proteins containing at least one GAF domain. From an evolutionary perspective, the GAF domain is a multi-purpose sensor found in all kingdoms of life. Therefore, it is unlikely that most of these proteins are CBCRs. Because we were interested in comparing nuances of CBCR GAF domain characteristics and their effects on chromotype, we needed to construct a working dataset of proteins, specifically enriched for photoactive GAFs. Automated database annotations appeared to be unreliable in distinguishing between CBCR and other GAF domain types.

We searched the literature for primary studies looking at one or more CBCR GAF domains. Domains that were either: 1) spectrally characterized with available data, or 2) manually analyzed and predicted to be CBCR in nature, were included in our dataset. Some publications/patent applications have quoted the spectral tune of certain GAF domains, but absorbance spectra were not available. In those cases, the domain was considered predicted rather than characterized.

Additional GAF domains were also included. It is common for two or more CBCR GAF domains to occur together on the same protein. We thus extended our dataset to neighboring GAF domains on the same proteins which we identified with InterPro. We also included putative GAF domains from the genome of *Synechococcus* sp. PCC 7002 due to our specific interest in this cyanobacterium. These last GAF domains were located with a BLAST-P search against the genome of the strain using the TePixJg amino acid sequence as template and with the help of InterPro.

Some duplicate entries existed due to references using different names for the same gene/species. These duplicates were removed from our dataset. Finally, we removed entries where the amino acid sequence was unobtainable. The result was 242 GAF domains that fit our criteria. These domains were located on 132 proteins out of 36 species (Table 1).

**Table 1:**
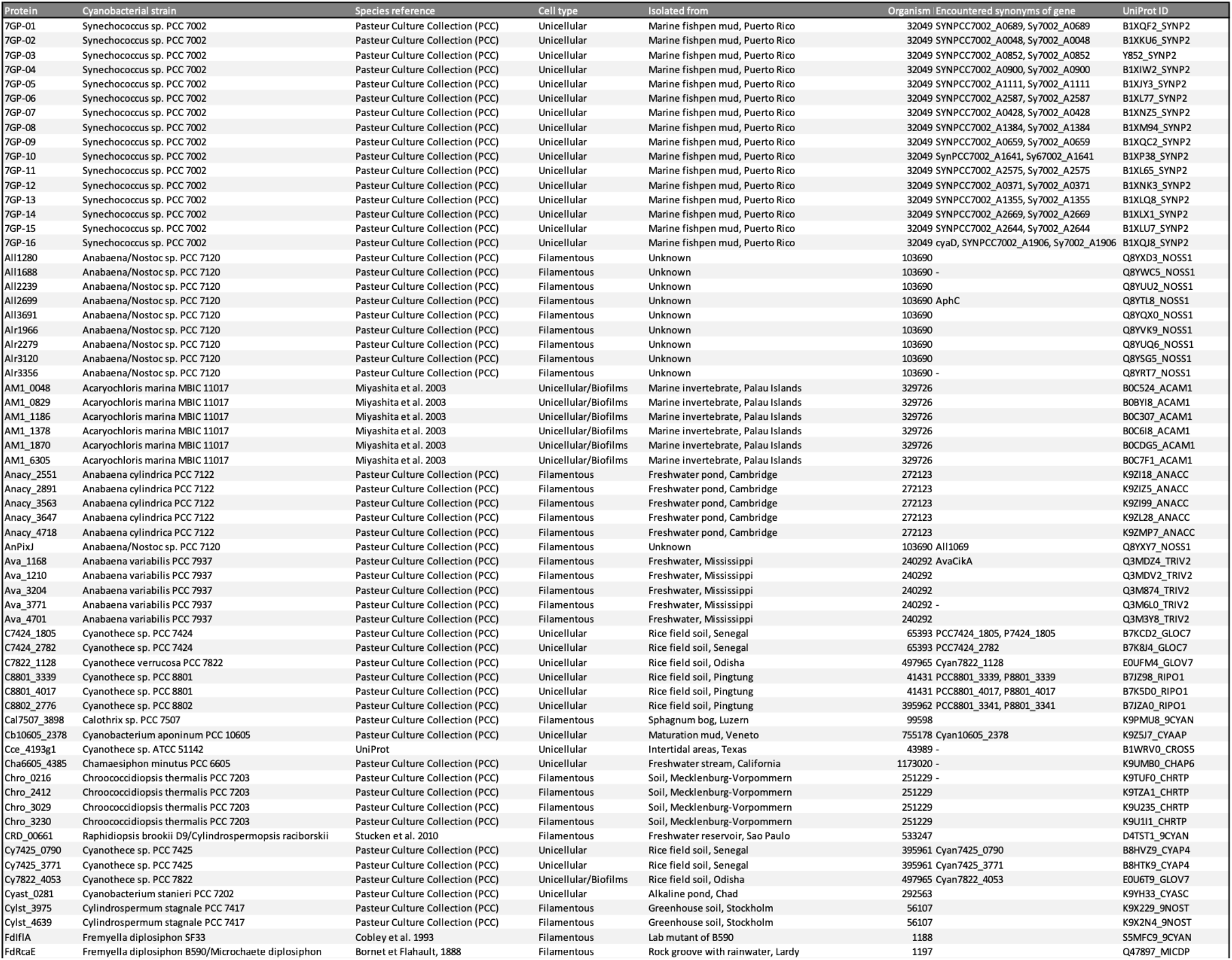
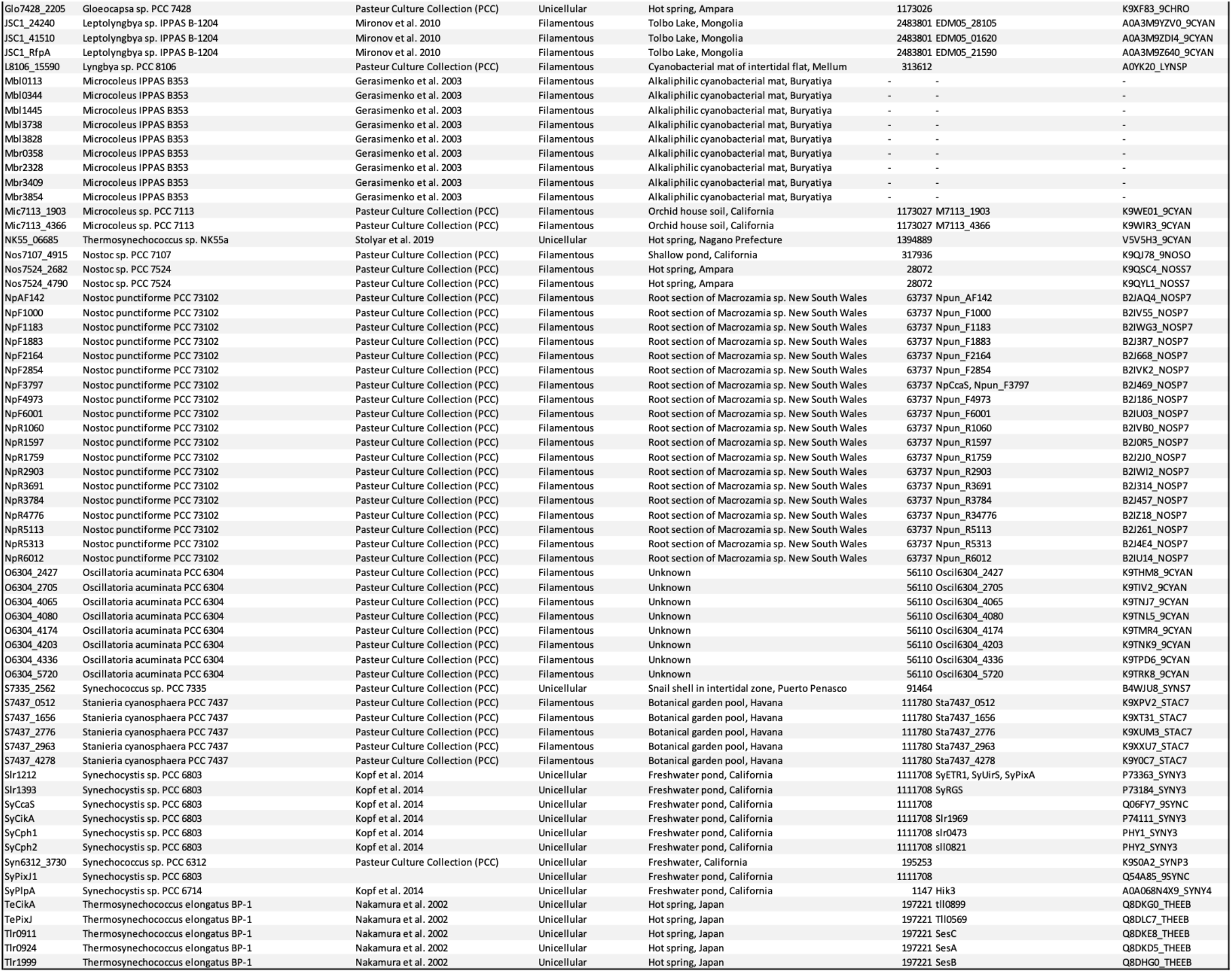
Metadata on species/proteins for the analyzed GAF domains.

98 of these GAF domains were spectrally characterized and formed the basis of a subset of entries. The λmax values of these spectra were used to convert the numerical spectral data into a set of categorical values for each GAF domain. This set of variables corresponded to chromotype as the colors absorbed by the 15Z/15E states. The following scale was used to assign λmax values to colors. Values were rounded to the nearest step of 50 nm and those above 700 nm are referred to as “Far-Red” (F). Similarly, values around: 450 nm ∈ “Blue” (B); 500 nm ∈ “Teal” (T); 550 nm ∈ “Green” (G); 600 nm ∈ “Orange” (O); 650 nm ∈ “Red” (R). These colors may differ from the ones quoted for GAF domains in the original publications as there does not appear to be a consensus in the literature regarding the relationship between spectral color and wavelength. We observed 19 different flavors of chromotype.

### CBCR GAF domain families

Our first hypothesis was that the overall sequence affiliation of GAF domains is a good predictor of chromotype. To do such an analysis would require first grouping GAF domains based on their present sequence similarity. The literature contains some well-known CBCR lineages/families. Examples include the Red-green, DXCF, DXCIP, Green-red, Insert-Cys, Dual-Cys, New Insert Cys, etc. Most of these names are informative about certain properties/features of a GAF domain but tend to be one-sided and difficult to assign readily. Therefore, the family (or lineage) of GAF domains was not recorded from the literature. Instead, the GAF domains were grouped phenetically using the Neighbour-joining algorithm and bootstrap value Figure 1.

**FIGURE 1:**
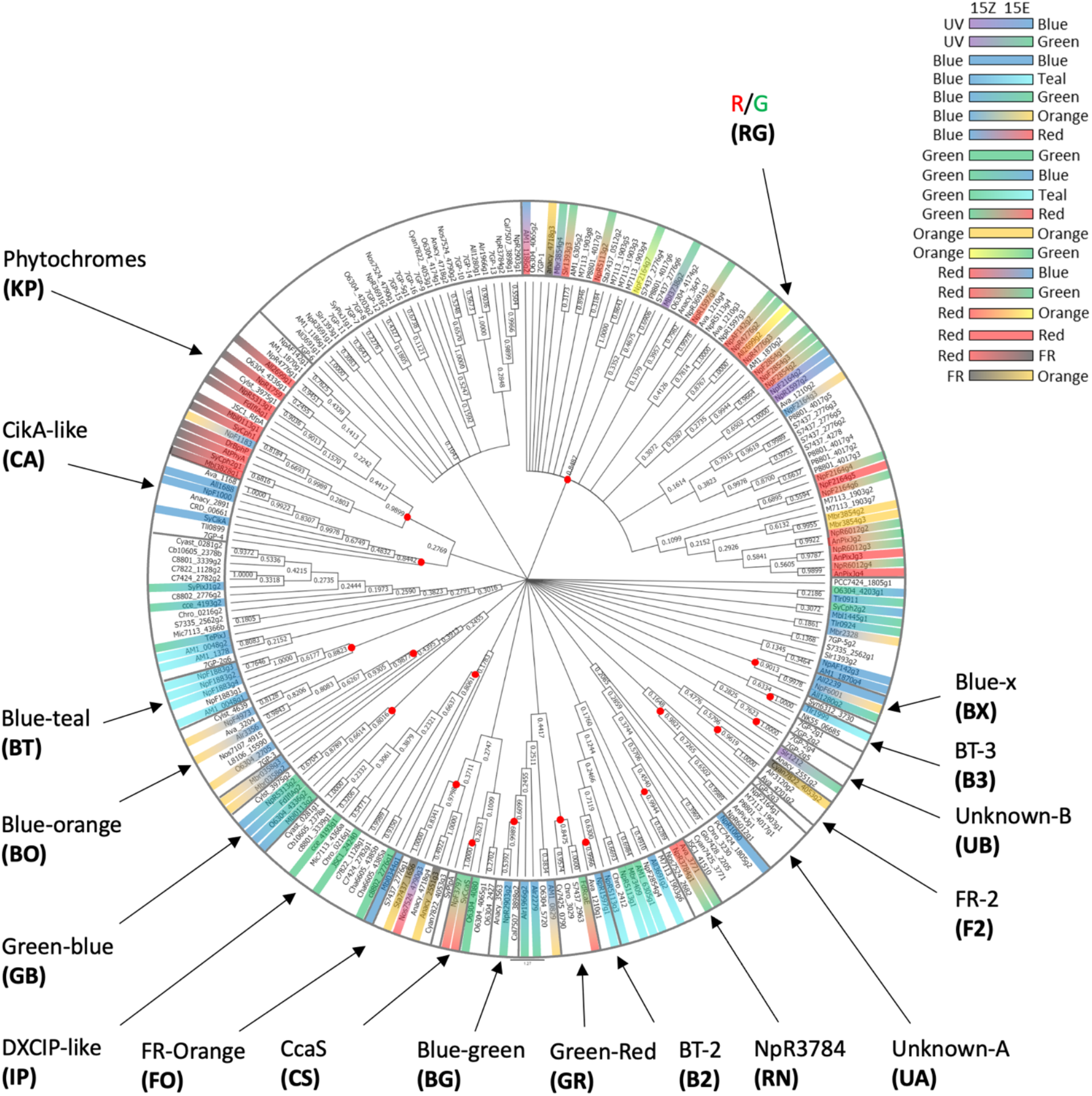
The GAF domain sequence is a good chromotype indicator. Data on GAF domains was collected and processed as described in the main text. The amino acid FASTA sequences of the 242 GAF domains were aligned using ClustalW. The sequences were assorted into clades using the Neighbour-Joining method. The evolutionary distances were computed using the Poisson correction method. Uniform evolutionary rates were assumed. All ambiguous positions were excluded from the comparison. The bootstrap consensus tree shown was inferred from 1000 replicates. The ratio of replicate trees in which the associated taxa clustered together are shown next to the branches. Branches reproduced in less than 10% bootstrap replicates are collapsed. The names of GAF domains are the same as the ones used in the literature. The absorbed colours, if known, are shown for both 15Z (inside) and 15E (outside) states. The GAF domains were then assigned to families, shown with arrows. Families were designated as the largest clades where associated taxa cluster together in at least 80% of the replicates. Families were named according to the most prominent known chromotype. The red dots indicate the beginning of those clades. Short names of each family used throughout this work are shown in brackets.

The result was 18 different families containing photosensitive GAF domains. Effort has been made in naming the families based on most prominent chromotype to make comparisons with the literature easier. However, using sequence motifs such as DXCF or “Insert Cys” to name families was avoided as much as possible. These tend to occur sporadically throughout various CBCR taxa and do not serve as reliable family identifiers. Some of the groups defined in this work correlate well with groups defined in the literature in terms of GAF domain members. For example, the RG group correlates well with the “Red-Green” family. Similarly, the BO group correlates with the “Blue-Orange” family, the BT group with the Blue-Teal family, the CA group with the CikA-like family, the IP group with the DXCIP family, the RN group with the NpR3784-like family. Some variation in the member domains was present.

In general, the overall sequence affiliation of a GAF domain can be a good general indicator of chromotype. It was obvious that a high sequence similarity generally translates to chromotype similarity. For example, only 7% of RG members do not absorb at least one colour at 550 nm or more. Only one member (9%) of the phytochrome family does not have the R/F chromotype. There were significantly less characterised members from the other groups. However, there was a clear indication of a conserved chromotype and spectral position. Examples are the BT, CA, BO, GB, IP and RN groups. About half of the CBCR GAF domain families have a strong preference for a chromotype. The rest of the families are not as well characterized. Suppl. Fig. 1 shows the distribution of chromotypes according to family.

Some chromotypes are more endemic whereas others are more sporadic. The B/O chromotype is currently the only known chromotype in the BO group. It is also present in the RG, RN and RX families as well as in some unclassified GAF domains. This ubiquity of B/O may be indicative of its usefulness and/or mechanistic simplicity. On the other hand, the most conservative chromotype is R/F, endemic to knotless phytochromes which is an outcome of knotless phytochromes using a different bilin (phycochromobilin, PФB). The R/G chromotype is the most abundant in our dataset yet almost endemic to the RG group. This is likely a reflection of the chromotype being almost exclusively found in the RG group which is oversampled in our dataset. R/G is interesting due to resembling the chromotype of free PCB in solution. We hypothesize R/G to be the “default” chromotype for relatively undifferentiated GAF domains. The distribution of R/G among families indicates that it is likely relatively easy to de-tune or not particularly useful. The RG group itself is the most spectrally diverse, suggesting that RG and R/G could provide a relatively quick way to adapt spectrally. In other words, the R/G chromotype has the shortest evolutionary distance to all other chromotypes. Finally, blue light absorption (as either 15Z or 15E) is the most common color perception, likely due to it requiring a single insert Cys residue in the binding pocket of the domain.

### CBCR domain architectures

The next hypothesis is that similar CBCR domain architectures will likely have similar chromotypes. We are not aware of a more systematic analysis of CBCR domain architectures at the time of writing. CBCR proteins tend to be multi-domain and often including more than one type of sensory domain, in addition to an effector domain. It is presently unknown how signal transduction from all these sensory domains is integrated to the effector domain. Therefore, we cannot assume that the order of the domains on the protein is important. This consideration implies that whole protein sequence alignments may not be the best way to characterize CBCR architectures. Instead, a more general and position-independent approach is to be used. The Principal Component Analysis (PCA) was chosen where the various types of domains are the input variables, their number in each protein is the weight component, and the CBCR proteins from the dataset are the individuals (Sup. Fig. 2). The results of the analysis are shown in Figure 2a while the basis of the Cartesian distribution is shown in Figure 2b. The results of the figure indicate the presence of at least four clearly defined CBCR domain architectures, summarized in Figure 3.

**Figure 2:**
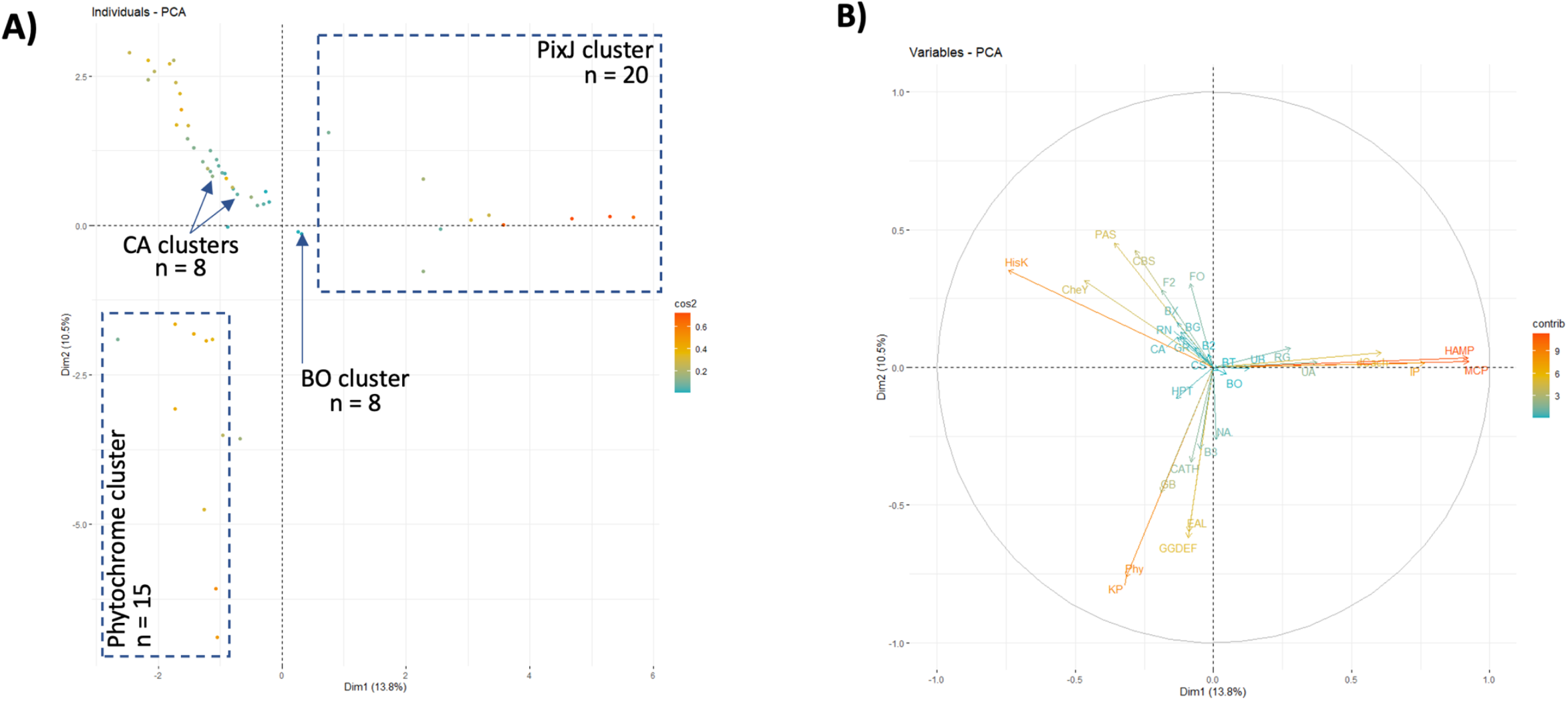
CBCR protein architectures can be defined with PCA. The number of each type of domain was counted for every CBCR listed in [Suppl. Fig 1 / Slide 15] and used as input for PCA. A) Position of individual genes with respect to the first two principal components, explaining 13.8% (PC1, x-axis) and 10.5% (PC2, y-axis) of variation, respectively. Genes are coloured according to the squared cosine. B) Map of the relationship between variables used in the PCA. Positively correlated variables point in the same direction, negatively correlated variables point in opposite directions. Arrow length and colour indicate the contribution to the principal components (blue: low, red: high). Domain types only found once were removed as categories from the input dataset. GAF domains without a designated family were also removed from the input dataset.

**Figure 3:**
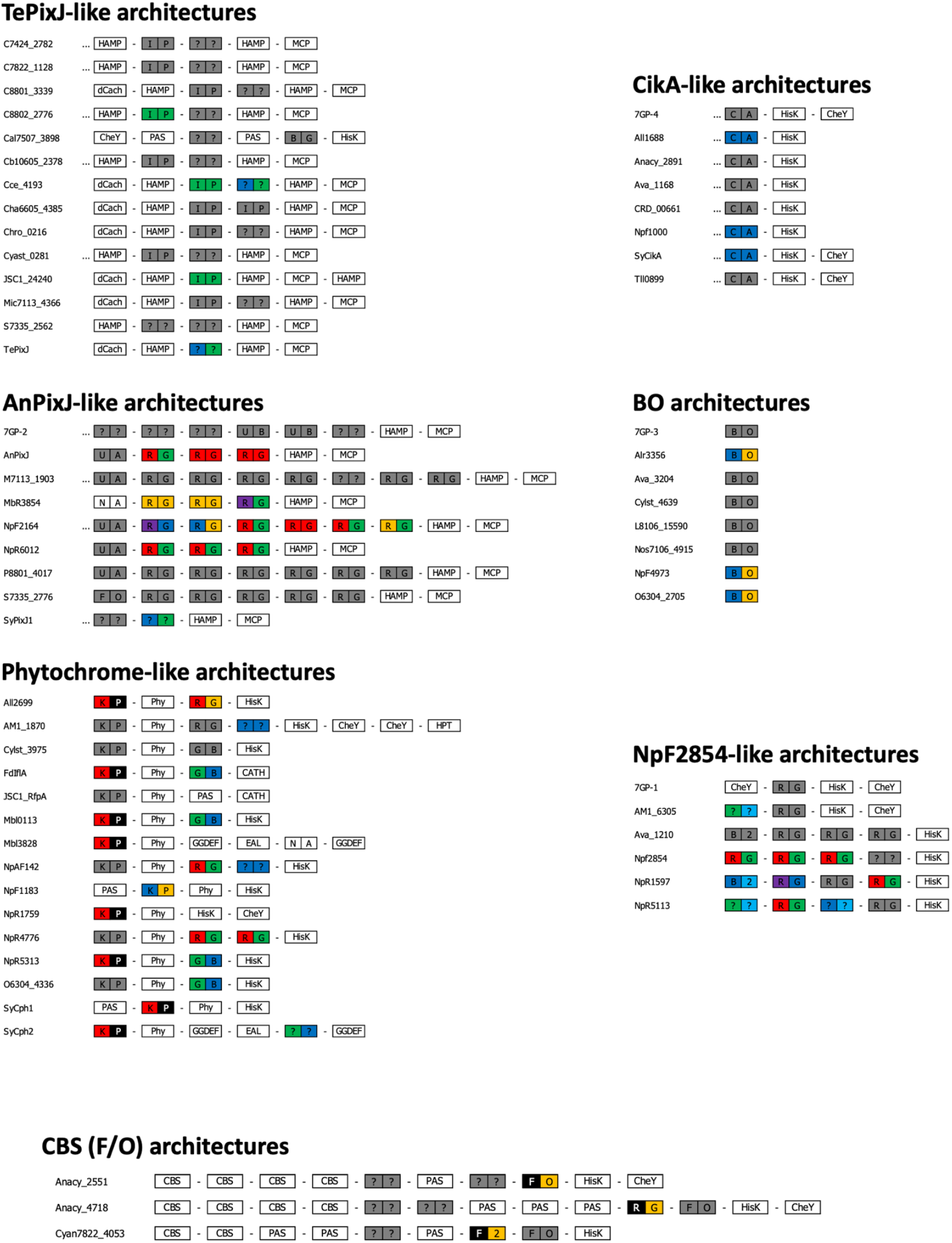
Types of CBCR protein architectures. For each of the genes listed, the predicted domain structure of the encoded protein is shown from N-terminus (left) to C-terminus (right). GAF domains are marked as rectangles with two boxes for 15Z (left) and 15E (right) states, filled according to the absorbed colour, if known. Unknown colours are marked in dark grey. Letters refer to the families assigned in Figure 2. GAF domains for which a sequence was unavailable are marked as “N/A”. Regions with unrecognized domain signature are marked as “…”. The naming of other domains follows InterPro convention.

#### Knotless phytochrome architecture

This type of architecture is primarily defined with the presence of a CBCR GAF domain from the KP family, always followed by a Phy domain. There is a strong tendency for the presence of a GB GAF domain and in other cases the RG one. Thus, the phytochrome architecture normally contains the [KP]-[Phy]-[GB]/[RG] domain motif. Interestingly, the effector domain portion is less constricted where the choice is between [GGDEF]-[EAL], a HisK, or a CATH domain. In nearly all cases this architecture focuses on R/F perception. There appears to be strong selection for an additional G/B chromotype, typically but not always provided by the GB GAF domain family. Where an RG domain is present it is usually with the more classical R/G chromotype.

#### AnPixJ- and TePixJ-like architectures

These two CBCR domain architectures are defined by the presence of a [HAMP]-[MCP] domain motif at the C-terminus. In the case of TePixJ-like architectures, there is an additional HAMP domain at the beginning of the N-terminus. Together these two HAMP domains flank one or two CBCR GAF domains, most commonly of the IP family. AnPixJ-like architectures also terminate in [HAMP]-[MCP] but this motif is preceded by a series of GAF domains, typically from the RG lineage. Currently, the AnPixJ architectures seem to be the most spectrally diverse, pertaining to their preference for RG domains. TePixJ architectures presently focus on green light perception.

#### CikA architectures

This type of CBCR proteins consists of a single GAF domain, followed by a HisK. The GAF domain is always from the CA group. The HisK is sometimes followed by a CheY domain. Currently, all known CA GAF domains are of the B/B chromotype.

#### BO architectures

These are the simplest CBCR architectures possible and consisting of only a single GAF domain. The GAF domain is of the BO family. Currently, the only observed chromotype in this family is the B/O.

#### NpF2854-like architectures

These architectures are reminiscent of the AnPixJ ones in incorporating several RG GAF domains. Although NpF2854 and AnPixJ architectures are separated in the Cartesian space, they could be thought of as subsets of the same architecture design. This is obvious from Sup. Fig. 3a where all RG-containing proteins are shown. Instead of a [HAMP]-[MCP] C-terminal portion, there is a HisK. The enzymatic HisK is sometimes followed by a CheY domain. Although this architecture is relatively unexplored, it does seem relatively chromatically diverse.

#### CBS architectures

The CBS architectures are characterized by the presence of two or more CBS domains on the N-terminus, several PAS domains in the middle and at least one GAF domain with the F/O chromotype towards the end. Currently, all three members have a HisK effector domain. What is interesting about this architecture is that the F/O GAF domains are of different families: FO, RG and F2, illustrating the convergent evolution of colour perception. There is an additional uncharacterized FO GAF domain in 2/3 members, with a high likelihood of an F/O chromotype. Currently, none of the other GAF domains have been characterized.

The various domain architectures have a strong preference for specific GAF domain families. It seems that the domain architecture could be a great supplemental chromotype predictor for cases where this is not certain or obvious from the GAF domain family per se. Currently, there’s only 18 putative proteins with more than one spectrally characterized GAF domain Sup. Fig. 3b. As more of these proteins become available, more certain predictions can be made based on domain architecture. In all cases, the architecture context (and spectral function) for each GAF domain could be indicated by its two neighboring domains.

### Structural diversity and chromotype

#### Most 3D variation is in the binding pocket

Chromotype is known to be a function of the shape of the binding pocket, in addition to the properties of some residues therein. We suspected that sequence alignments per se would not be sensitive to nuances in amino acid sequence which yield similar 3D shapes (and affect chromotype). To go around this problem and provide a basis for even more accurate chromotype predictions based on available data, we simulated the 3D structures of all 98 characterized GAF domains and constructed a contact map (Figure 4). The results of our analysis show that the majority of structural variation between the predicted GAF structures is located in the chromophore binding pocket. This observation is in line with our expectations given the diversity of chromotypes among known CBCR GAF domains.

**Figure 4:**
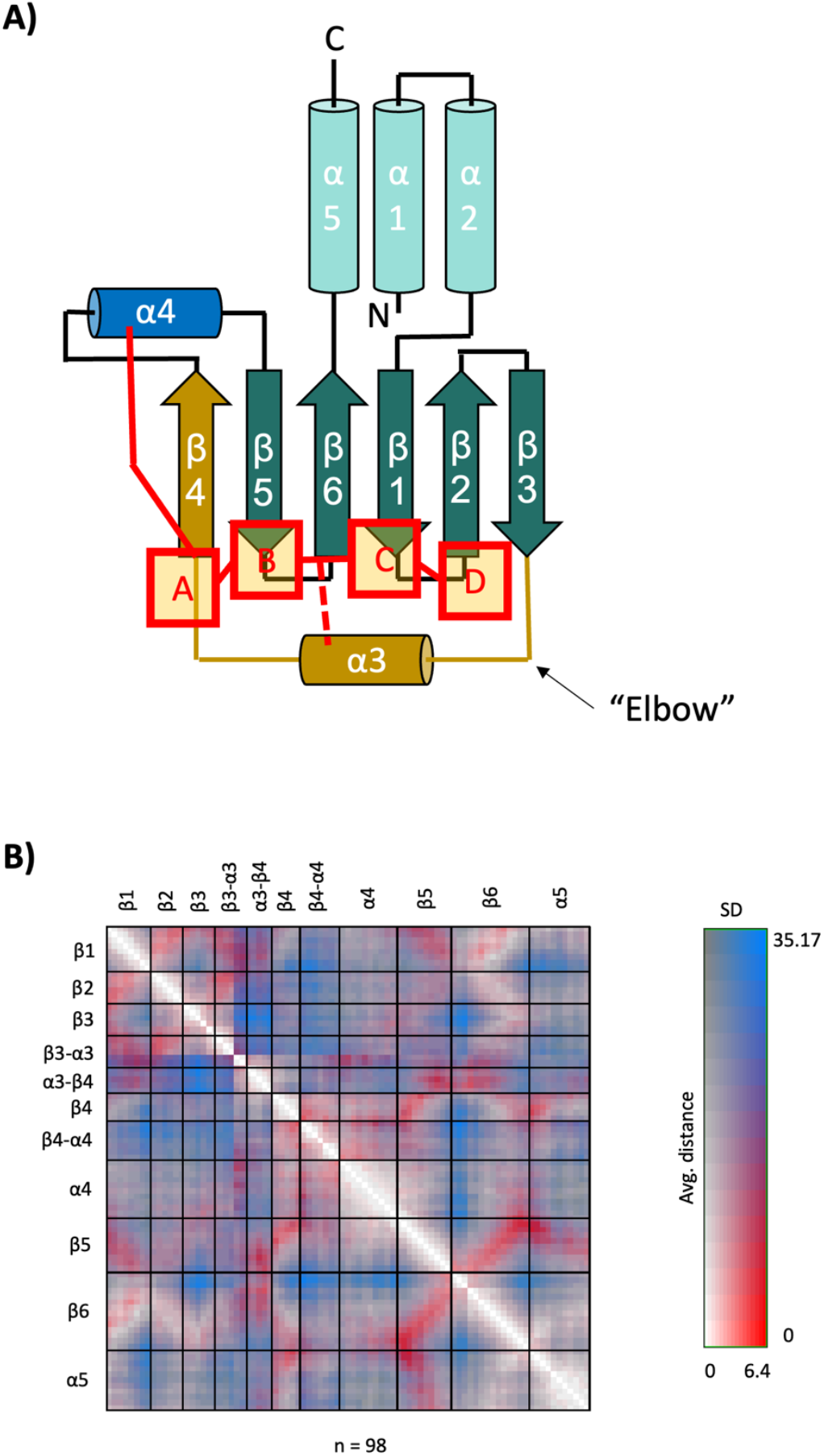
Most structural variation is within the chromophore-binding pocket. The three-dimensional crystal structure of TePixJ (3VV4) was used as a template to create a topology model of the CBCR GAF domain. A) The GAF domain contains a central planar region of five anti-parallel β-sheets (β1,2 and β4-6). The N- and C-termini are located on one side of the central region and form a bundle of three α-helices (N-terminal α1,2 and C-terminal α5). The region between β3 and α4 (shown in gold) is variable between photoactive GAF domains. Helix α4 (blue) contains a conserved Cys residue in all families except DXCIP (IP). This residue binds the bilin chromophore (red) non-reversibly at ring A. A Second Cys residue often occurs between β4 and β3. This residue can bind the C10 of the chromophore, between rings B and C. B) The three-dimensional structure of 98 GAF domains with a known chromotype was predicted using Phyre2 (Kelley et al. 2015). The residue-residue (RR) distance map shown here was calculated using RRDistMaps (Chen et al. 2015) and shown here. The map shows the positions of all Cα relative to each other. The map uses a two-dimensional scale shown on the right. Small average distances and small SD are shown in white. Small average distances and large SD are shown in red. Large average distances and small SD are shown in dark grey. Large average distances and large SD are shown in blue.

#### Grouping GAFs based on shape

Large-scale structural predictions enable high-precision/high-throughput comparative studies of structures. This is partially because of the great reproducibility of the method. At the time of writing there were only a handful of crystal or NMR structures available for CBCRs. We decided to split the consensus structure of a typical CBCR GAF domain (Figure 4a) into 14 categories, corresponding to different parts of the structure. We aligned the 3D structures of the GAF domains and compared all 98 structures at each of the 14 regions. If the same region looked identical between two or more GAF domains, they were assigned the same (non amino-acid) letter, from A-Z. The result is a string of 14 letters that perfectly describe the 3D structure of a GAF domain, relative to others. Supplementary Figure 4 shows contact maps of four groups of GAF domains with identical structural strings. Differences in 3D structure were confined to only one or two individual residues.

Inspired by linguistics, we decided to calculate the Levensthein distances between all structural strings to allow for a form of sequence alignment, similar to DNA/protein sequences but directly relating to 3D shape instead. The results are shown in Figure 5a. Three large structural families were identified from this analysis which we refer to as “IBDC”, “ACAB”, “AAEI”. The colors of light absorbed by members of each of these families, irrespective of 15Z/15E, were counted and represented as a percentage of all members (Figure 5b). The raw data is available in Sup. Figure 5. Blue light absorption is thought to be achieved by means of a single residue: an insert Cys in the binding pocket. If blue light absorption is ignored, the results show that the three structural groups specialize in three broad regions of the spectrum. AAEI is preferred for absorption from green (550 nm) to far-red (700+ nm). ACAB is more suited from green (550 nm) to red (650 nm). IBDC seems preferable from teal (450 nm) to green (550 nm). The general 3D shapes of the GAF domains could thus be thought of as a coarse control for the spectral sensitivity of the domain.

**Figure 5:**
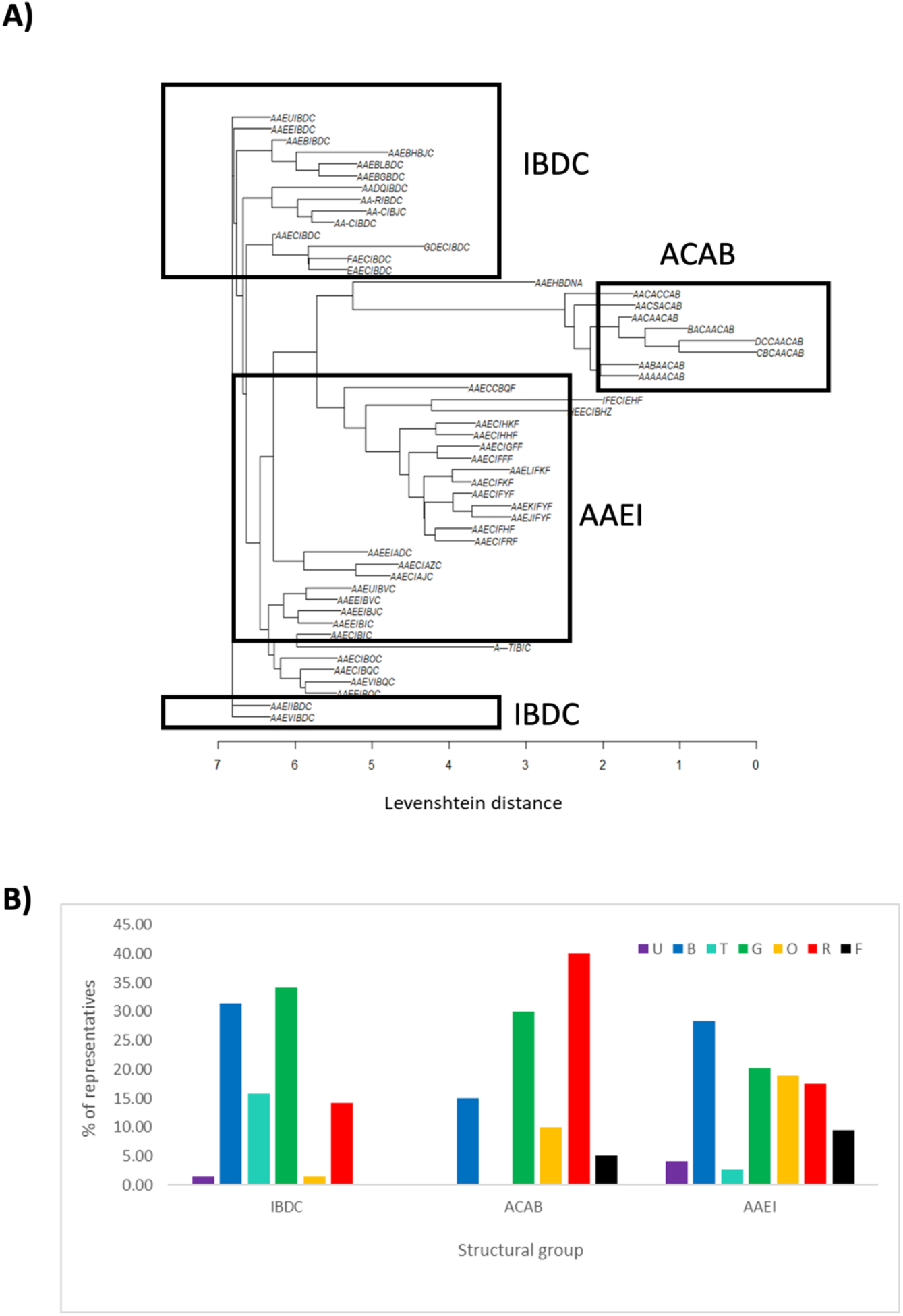
Grouping of GAF domains based on predicted structure reveals link to chromotype. The strings describing the binding pockets 3D shapes of all 98 characterized GAF domains were collected. These include: β1-β2, β2-β3, β3, “elbow”, β3-α3, α3-β4, β4, β4-α4 and α4. The Levenshtein distances between all 52 unique strings were calculated using a custom algorithm. A) These distances were used to construct a guide tree using the Neighbour-Joining algorithm. The length of the branches corresponds to the Levenshtein distance. The IBDCAAEI-, and ACAB-type groups are labelled. B) The number of states (both 15Z and 15E) absorbing a given colour were counted for each GAF domain with a given structural configuration. These were then represented as a frequency based on the total number of GAF domains with said configuration.

### Key binding pocket residues

Fushimi et al. (2020) identified six key positions in the CBCR binding pocket influencing chromotype. One of these key residues is the insert Cys, mentioned above. The key residues for each of the six positions were identified in all 98 characterized domains from our dataset.

### Chromotype predictions

Supplementary Figure 6 contains a summary of all insights for the 98 characterized GAF domains acquired from this study. Individually, none of the four characteristics of CBCR diversity (GAF family, protein architecture, 3D structures, key residues) matched well with chromotype during preliminary studies. We hypothesized that the current dataset size will be impractical for machine learning approaches and instead elected to use another dimensionality reduction technique but for categorical data: the Multiple Correlation Analysis (MCA). The results of this diversity analysis are shown in Figure 6. The results do show the expected continuity of characteristics, but some clear clusters could be identified on the Cartesian space. Moreover, there is not a significant overlap of the confidence ellipses, enabling us to use them as an indicator of the two most likely colors (15Z and 15E) absorbed by an unknown GAF domain.

**Figure 6:**
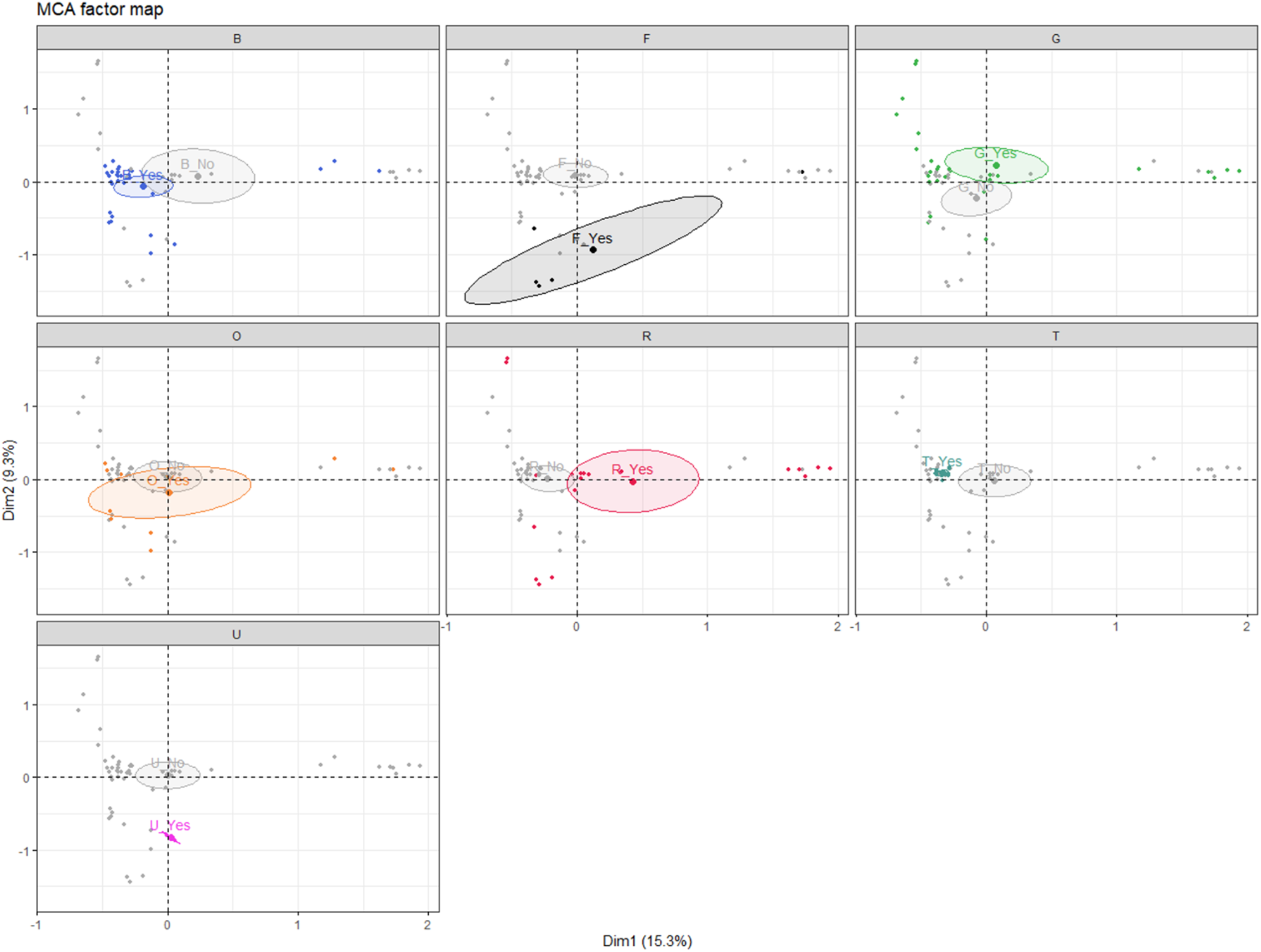
Summary of characterized GAF domain characteristics. The structural string configuration, in addition to the binding-pocket residues were gathered for all 98 characterized GAF domains. These variables were used as input for an MCA. The absorbed colours (irrespective of 15Z/15E) were used as supplementary variables which do not affect the clustering. A: Position of individual genes with respect of the first two principal components, explaining 15.6% (PC1, x-axis) and 8.8% (PC2, y-axis) of variation. Each panel shows the same dataset according to absorbed colour. “Yes” (labelled at the centre of each confidence ellipse) indicates that the individuals absorb the given colour. Likewise, “No” indicates that the individuals do not absorb the given colour. The ellipses in each panel reflect the 95% confidence region, according to colour.

#### GAF domains from *Synechococcus* sp. PCC 7002

The GAF domains from *Synechococcus* sp. PCC 7002 were subjected to the same analysis and the results are presented in Supplementary Figure 7. Only five of the 22 GAF domains (7GP-01g, 7GP-02g4, 7GP-02g5, 7GP-03g, 7GP-04g) could be mapped to existing GAF domain families and structural strings. The remainder of domains were outside of the confidence range established by this work. These GAF domains were used as supplementary individuals in the MCA analysis. 7GP-01 was within the confidence of prediction. Aside from the unique key residue six being Asp, 7GP-03 was also within the confidence of prediction. None of the other GAF domains from *Synechococcus* sp. PCC 7002 were good candidates for prediction. 7GP-01 was mapped as a supplementary individual to the MCA analysis and the result is shown in Figure 7a. We then recalculated the MCA analysis without key residue six in order to incorporate 7GP-03 too (Figure 7b).

**Figure 7:**
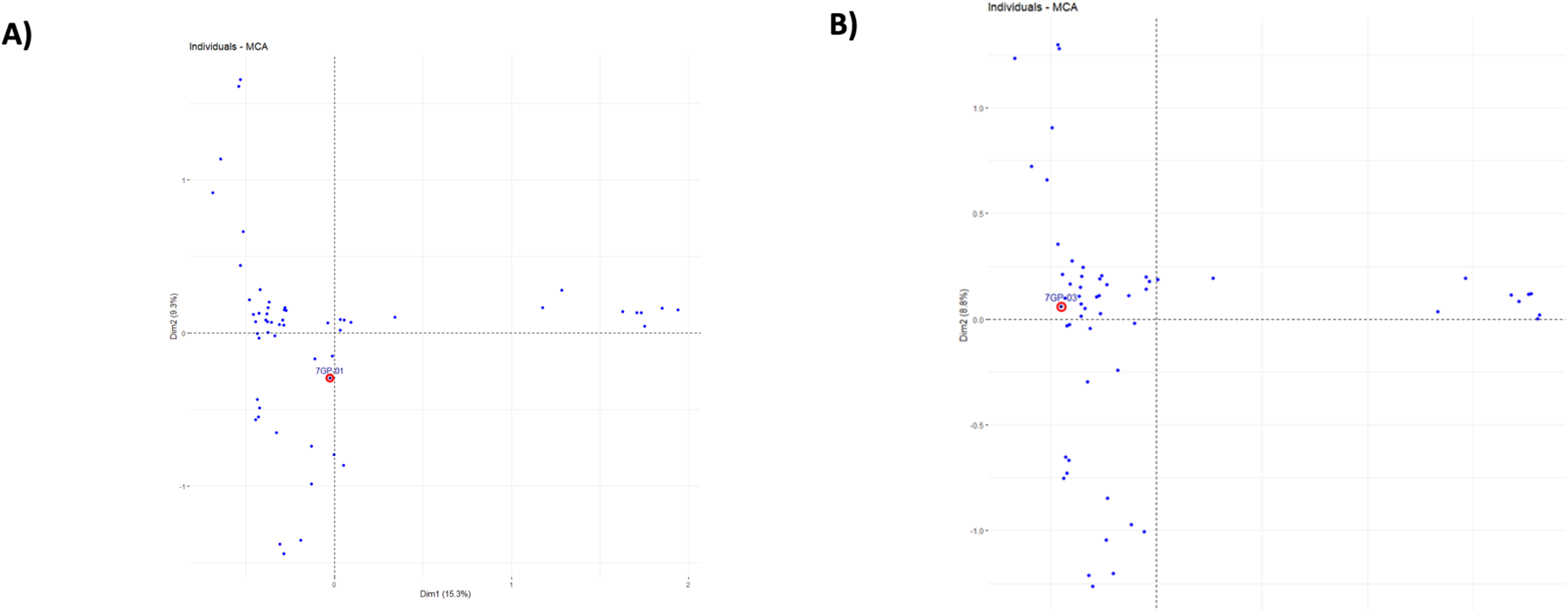
Chromotype predictions for 7GP-01 and 7GP-03. A) The predicted structure and sequence features of 7GP-01 were added to the results dataset from Figure 6. 7GP-01 was used as a supplementary individual which did not affect clustering. Position of individual genes with respect of the first two principal components, explaining 15.3% (PC1, x-axis) and 9.3% (PC2, y-axis) of variation. The clustering of 7GP-01 is shown in red. B) The predicted structure and sequence features of 7GP-03 were added to the results dataset from Figure 6. 7GP-03 was used as a supplementary individual which did not affect clustering. Position of individual genes with respect of the first two principal components, explaining 15.6% (PC1, x-axis) and 8.8% (PC2, y-axis) of variation. The clustering of 7GP-03 is shown in red

#### 7GP-01

Based on the MCA analysis there is 95% confidence that 7GP-01 is sensitive to orange (600 nm) and a 95% chance it is sensitive to red light. 7GP-01 is a member of the RG group but contains an insert Cys residue in the binding pocket, albeit at an unusual position likely away from C10 of the chromophore.

#### 7GP-03

The analysis indicates that there is a 95% confidence that 7GP-03 is sensitive to orange (600 nm) and a 95% chance it is sensitive to blue light. 7GP-03 is also a member of the BO family.

## DISCUSSION

Our review on the diversity of CBCR GAF domains showed that the sequence similarity generally translates to chromotype similarity. There are 18 families of CBCR GAF domains defined in this analysis. The groups occurred within similar CBCR architectures when clustered by 80% identity. In addition, some groups had stringent chromotypes at 80% similarity (BT, BO). Other groups had more varied chromotypes (RG). This means that the sequence similarity in the former groups is centered on the chromophore binding pocket. In the same way, the sequence similarity of the latter group is weighted towards different regions. The last conclusion is supported by the fact that RG GAF domains have a stronger preference to occur as GAF arrays, rather than being exclusive to one protein architecture.

It could be said that the binding pocket of RG domains is less evolutionarily differentiated (less defined). This is supported by Figure 6. The same figure shows that individuals sensitive to green, red, and orange light (RG members) are more spread out. In contrast, individuals sensitive to far-red, teal and UV-light form compact clusters. GAF domains that absorb blue light are an exception. Blue light sensitivity can be achieved with a single residue substitution. Because RG chromophore pockets are less differentiated, they should be spectrally more flexible. This has been well demonstrated by Fushimi et al. (2020). The absorbance spectra of RG domains are reminiscent of free PCB. This implies relatively less PCB compression is present at longer λmax values. This has been confirmed by NMR done by Cornilescu et al. (2014).

The spectral plasticity of RG domains corresponds to more chromatic diversity within similar architectures. RG domains co-occur as arrays within two architectures. These architectures are represented by AnPixJ and NpF2854 in Figure 3. The NpF2854-like architectures tend to be more homochromatic, whereas AnPixJ-like are more heterochromatic. Both architectures are more chromatically variable than others such as TePixJ-like. More chromatic plasticity is probably beneficial in adapting to more specialized environmental niches.

Evolutionarily differentiated groups absorb teal, far-red, UV and blue light. Examples are the BT, KP and CA groups. These appear to be preferred in architectures with less chromatic variability. Such architectures are likely to participate in more fundamental processes. CikA belongs to the CA group. This protein is a known clock modulator (Narikawa et al., 2008). Cph2 is known to elicit response to shading (Rockwell and Lagarias, 2017). Certain BT GAF domains are associated with cell aggregation (Enomoto et al., 2015).

An interesting case is the BO group. Nothing is known about its function. Members are highly conserved in chromotype and architecture. This suggests an essential role. A supporting observation from the literature is the case of Prochlorothrix hollandica PCC 9006 (Pinevich et al., 2012; Rockwell et al., 2015b). This cyanobacterium does not have phycobiliproteins. Moreover, the only recognized CBCR from this strain belongs to the Blue/Orange group.

There was an obvious preference for specific chromotypes in more specialized architectures. This preference could be used to fine-tune any chromotype prediction with low confidence. Our analysis on the predicted 3D shape of GAF domains also presents a way to predict chromotype, in addition to forming a hypothesis for further scrutiny.

According to our model, the overall spectral range of CBCRs is evolved via spacing restrictions of the binding pocket (“coarse focus”). Such restrictions are due to the predicted configuration of secondary structure elements within the binding pocket. These appear to be quite modular between GAF domains. Finer focus of the spectrum is achieved with smaller amino acid substitutions as proposed by Fushimi et al. (2020). The co-occurrence of elements from both the coarse and fine focus explains the majority of photocycles from the dataset.

We suspect that specific protein architectures are selected to satisfy certain spectral demands. The GAF domain family is thus a function of the various amino acid sequence biases, required by the protein architectures to achieve the desired chromotype. This was supported by our observation on CBS architectures having the same chromotypes but coming from different GAF families.

Overall, our review on CBCR diversity allowed us to make predictions on the spectral sensitivity of *Synechococcus* sp. PCC 7002. Further work can confirm these predictions and even extend them to other commercially interesting strains.

## METHODS

### ClustalW

Multiple sequence alignments were done with ClustalW. The gap opening penalty was set to 10, while the gap extension penalty was 0.2. The Gonnet weight matrix was used, where residue specific penalties were applied. Hydrophilic penalties were also applied. The gap separation matrix was set to 4, but without end gap separation. Finally, negative matrices were not used. The delay divergent cut-off was set to 30%. The predefined gap was not kept.

### Principal Component Analysis

The Principal Component Analysis (PCA) was done in R (4.0.2). The factoextra package was used. The .pca variables were computed using the prcomp() function. The graphs of individuals were computed using the fviz_pca_ind() function. The graphs of variables were computed using the fviz_pca_var() function. The raw data files for each experiment are attached in Appendix.

### Multiple Correspondence Analysis

The Multiple Correlation Analysis (MCA) was done in R (4.0.2). The FactoMineR and factoextra packages were used. The .mca variables were computed using the MCA() function. The graphs of individuals were computed using the fviz_mca_ind() function. The graphs of variables were computed using the fviz_mca_var() function. The raw data files for each experiment are attached in Appendix

## Supporting information

Raw output for generating Figure 1

All data

Python script to calculate Levenstein distances

Raw output for generating Figure 5

FASTAS of GAF domains

Aligned Fastas for Figure 1. Rename to .nwk

3D models

## SUPPLEMENTARY

Supplementary Figure 1 shows the chromotype distribution per family from Figure 1.

Supplementary Figure 2 shows the predicted domain architecture of all 133 proteins in this analysis, used for Figure 2.

Supplementary Figure 3 shows the domain architectures of proteins containing RG domains, as well as proteins with more than one characterized GAF domain (in aide to Figure 3).

Supplementary Figure 4 shows an analysis of the stringency of GAF domain grouping based on shape, used in Figure 4.

Supplementary Figure 5 shows the spectral colors absorbed by each GAF domain shape, used in Figure 5.

Supplementary Figure 6 shows the insights obtained for all 98 spectrally characterized GAF domains, used as an input for Figure 6.

Supplementary Figure 7 shows the insights obtained for the putative GAF domains from *Synechococcus* sp. PCC 7002, used for the predictions in Figure 7.

The Levenshtein distance calculations are done with a custom Python script, available as a supplementary file.

**SUPPL. FIGURE 1.**
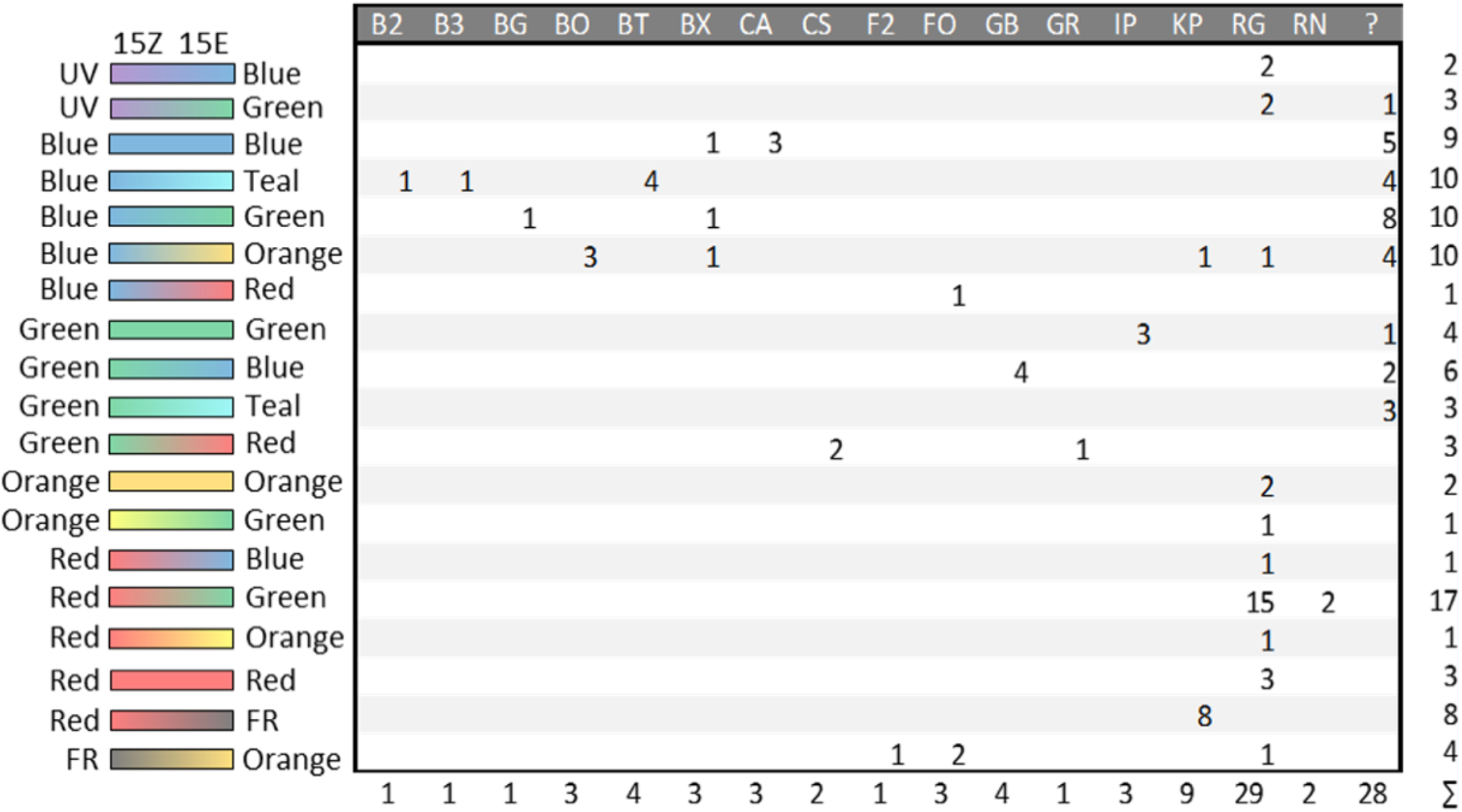
Chromotype distribution according to GAF family. The sums of GAF domains with the respective chromotypes or from the respective families are shown. Each two-letter code in the first row relates to a family assigned in Figure 3-2. There are no sub-families. Numbers indicate the amount of individual GAF domains (from 98 sequences) with characterized chromotypes listed in Figure 3-2. All characterized GAF domains which do not cluster into families are shown under the “?” column.

**SUPPL. FIGURE 2:**
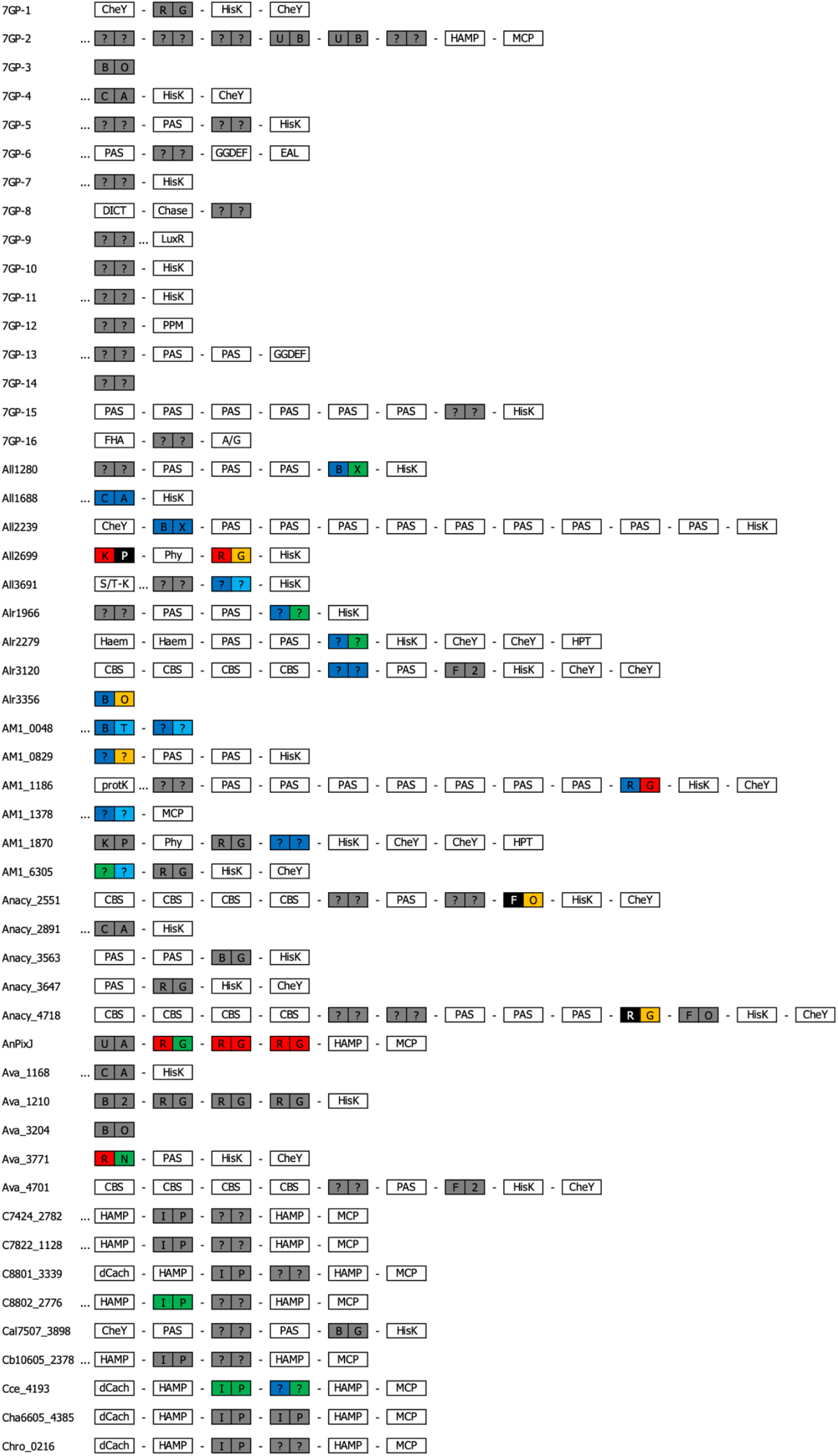
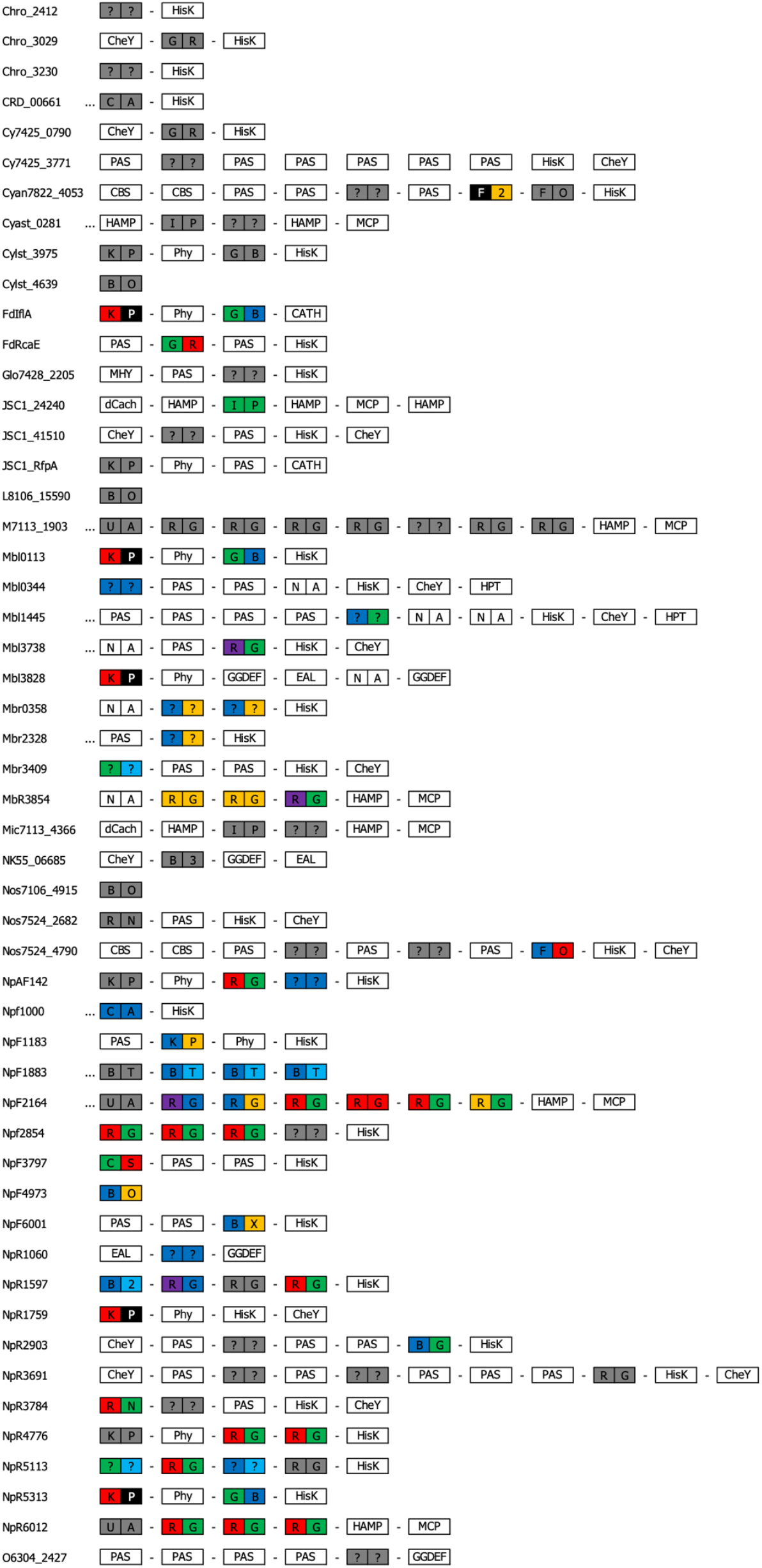
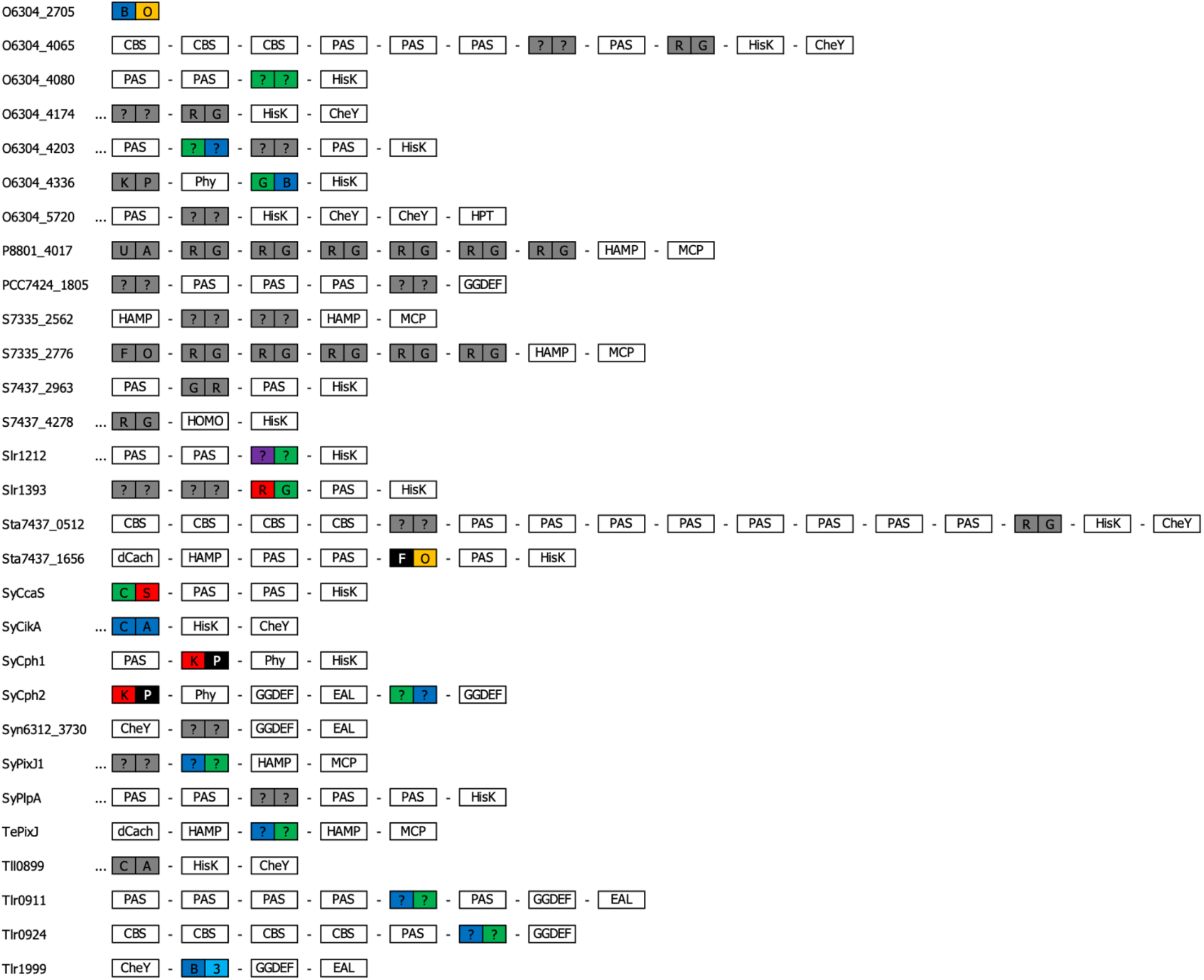
Domain architecture of CBCRs and GAF-domain proteins. For each of the genes listed, the predicted domain structure of the encoded protein is shown from N-terminus (left) to C-terminus (right). GAF domains are marked as rectangles with two boxes for 15Z (left) and 15E (right) states, filled according to the absorbed colour, if experimentally verified. Unknown colours are marked in dark grey. Letters refer to the families assigned in Figure 3.1. GAF domains for which a sequence was unavailable are marked as “N/A”. Regions with unrecognized domain signature are marked as “…”. The naming of other domains follows InterPro convention. A/G: Adenylyl cyclase class-3/4/guanylyl cyclase domain. dCach: dCache_1 domain. HOMO: Homodimerization domain. MHY: MHYT domain. S/T-K: Serine/Threonine Kinase domain.

**Suppl. Fig. 3:**
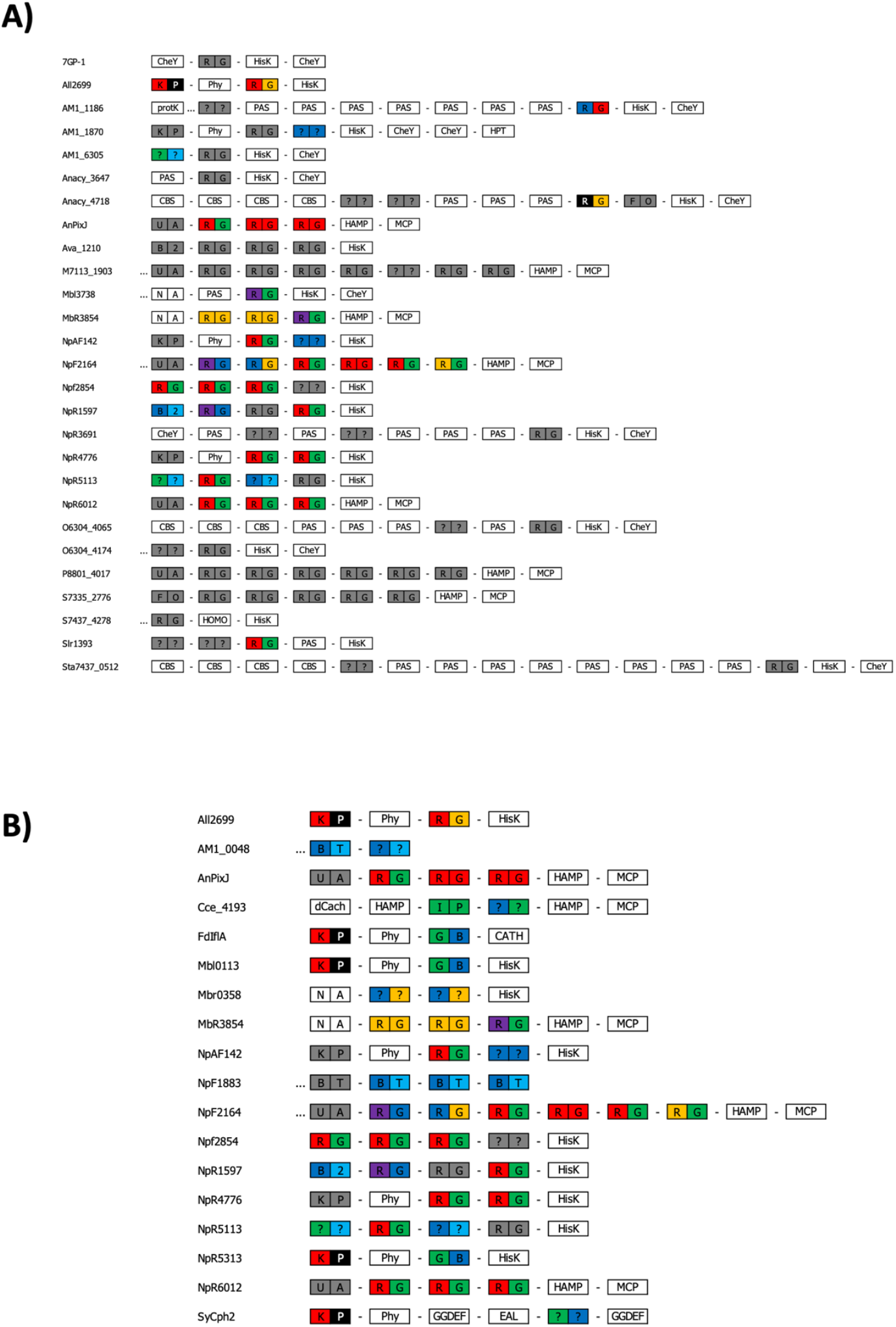
Demands imposed on CBCR properties. All CBCRs containing RG GAF domains are also shown (A). All CBCRs from this analysis that contain at least two spectrally characterized GAF domains are shown (B). For each of the genes listed, the predicted domain structure of the encoded protein is shown from N-terminus (left) to C-terminus (right). GAF domains are marked as rectangles with two boxes for 15Z (left) and 15E (right) states, filled according to the absorbed colour, if known. Unknown colours are marked in dark grey. Letters refer to the families assigned in Figure 3-2. GAF domains for which a sequence was unavailable are marked as “N/A”. Regions with unrecognized domain signature are marked as “…”. The naming of other domains follows InterPro convention.

**SUPPL. FIGURE 4:**
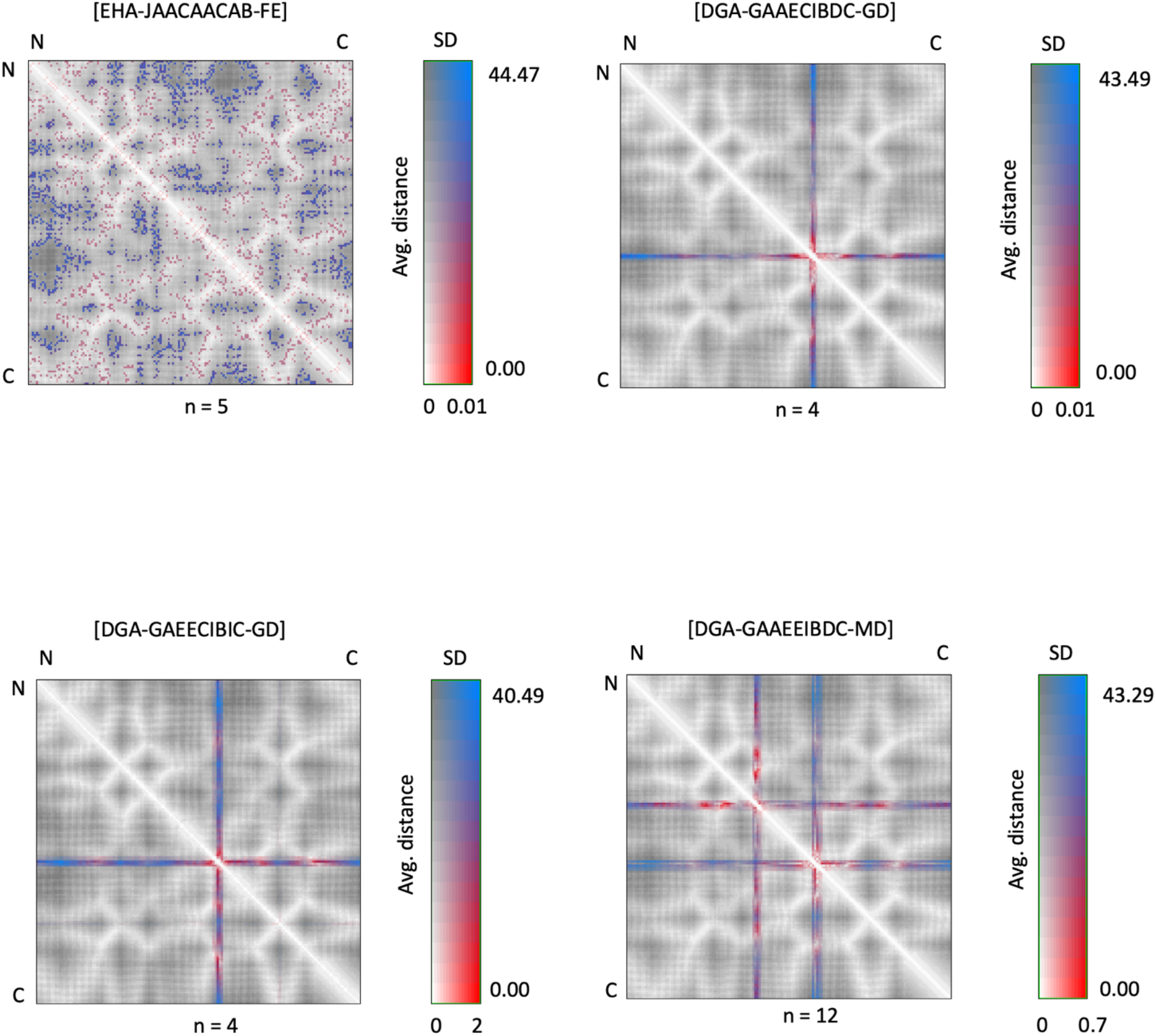
Categorization of predicted secondary structures of photoactive GAF domains. The three-dimensional structure of all characterized GAF domains was predicted using Phyre2 (Kelley et al. 2015). The structures were then superimposed with each other in the most optimal way using MatchMaker (Meng et al. 2006). The models were divided into regions of secondary structures. Each of those secondary structures was manually assorted into groups for every GAF domain. All secondary structures in the same position which have near-perfect overlap, length and overall shape share the same (non-amino acid) letter. The predicted structure of each GAF domain was thus summarized with a string of 14 letters (refer to Figure 3-26). In order: α1, α2, α2-β1, β1-β2, β2-β3, β3, “elbow”, β3-α3, α3-β4, β4, β4-α4, α4, β6-α5 and α5. The following regions were ignored: β1, β2, α4-β5, β5, β5-β6 and β6. GAF domains were then grouped according to matching string. The residue-residue (RR) distance map of all groups with at least 3 GAF domains was calculated using RRDistMaps and shown here. The map shows the positions of all α-Carbons relative to each other (refer to Figure 3-27). The map uses a two-dimensional scale shown on the right. Small average distances and small SD are shown in white. Small average distances and large SD are shown in red. Large average distances and small SD are shown in dark grey. Large average distances and large SD are shown in blue. The predicted string is shown above each panel. The number of GAF domains with this string are shown below the panel.

**SUPPL. FIGURE 5:**
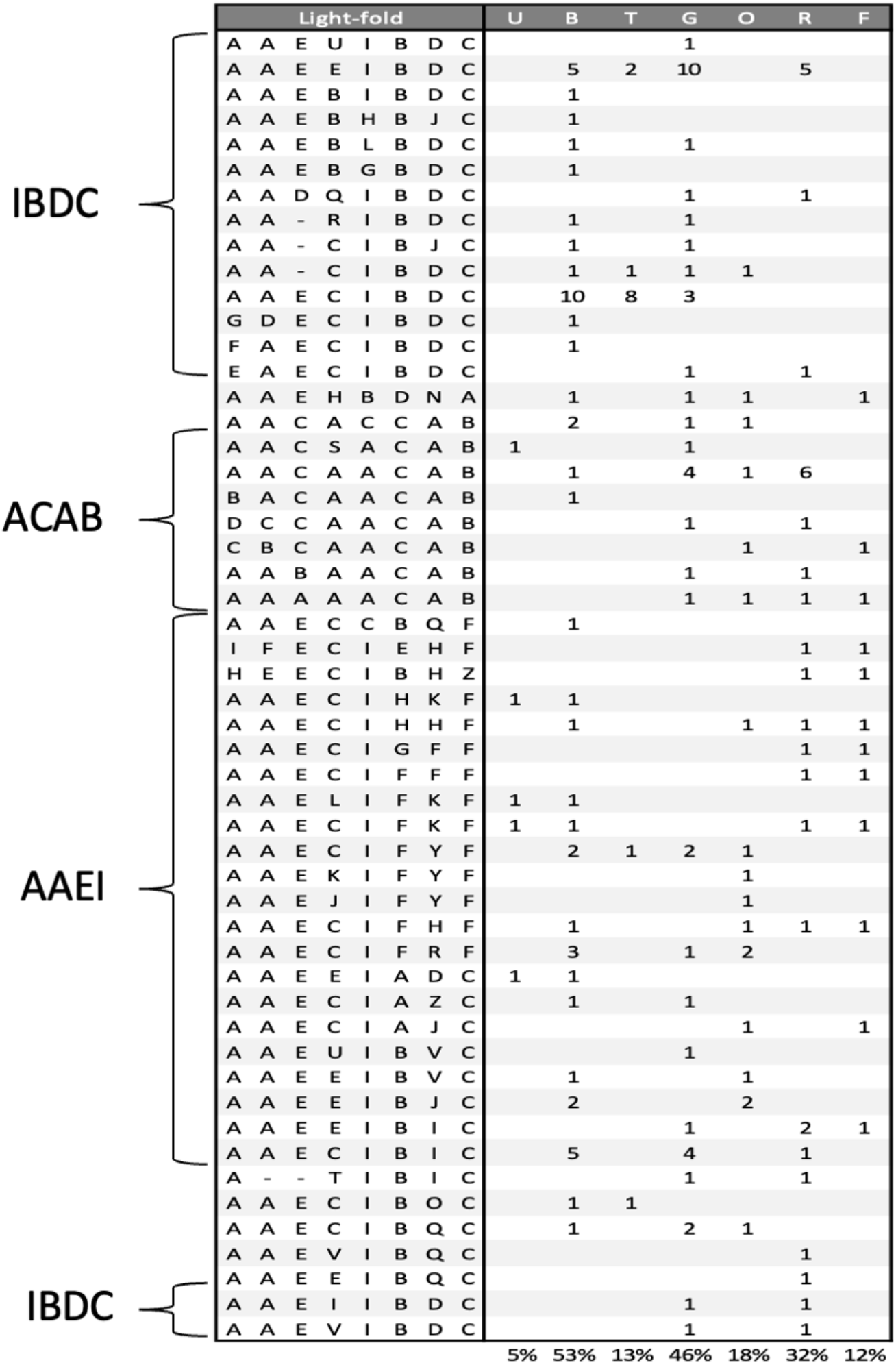
Spectral colours absorbed by each GAF structure type. The most readily absorbed colours for all 52 unique strings are shown on the right. The numbers show the number of GAF domains that absorb the relevant colour irrespective of 15Z or 15E. GAF domains which absorb the same colour in both isomers are counted once. The percentages below refer to the proportion of unique GAF domains that absorb a given colour.

**SUPPL. FIGURE 6:**
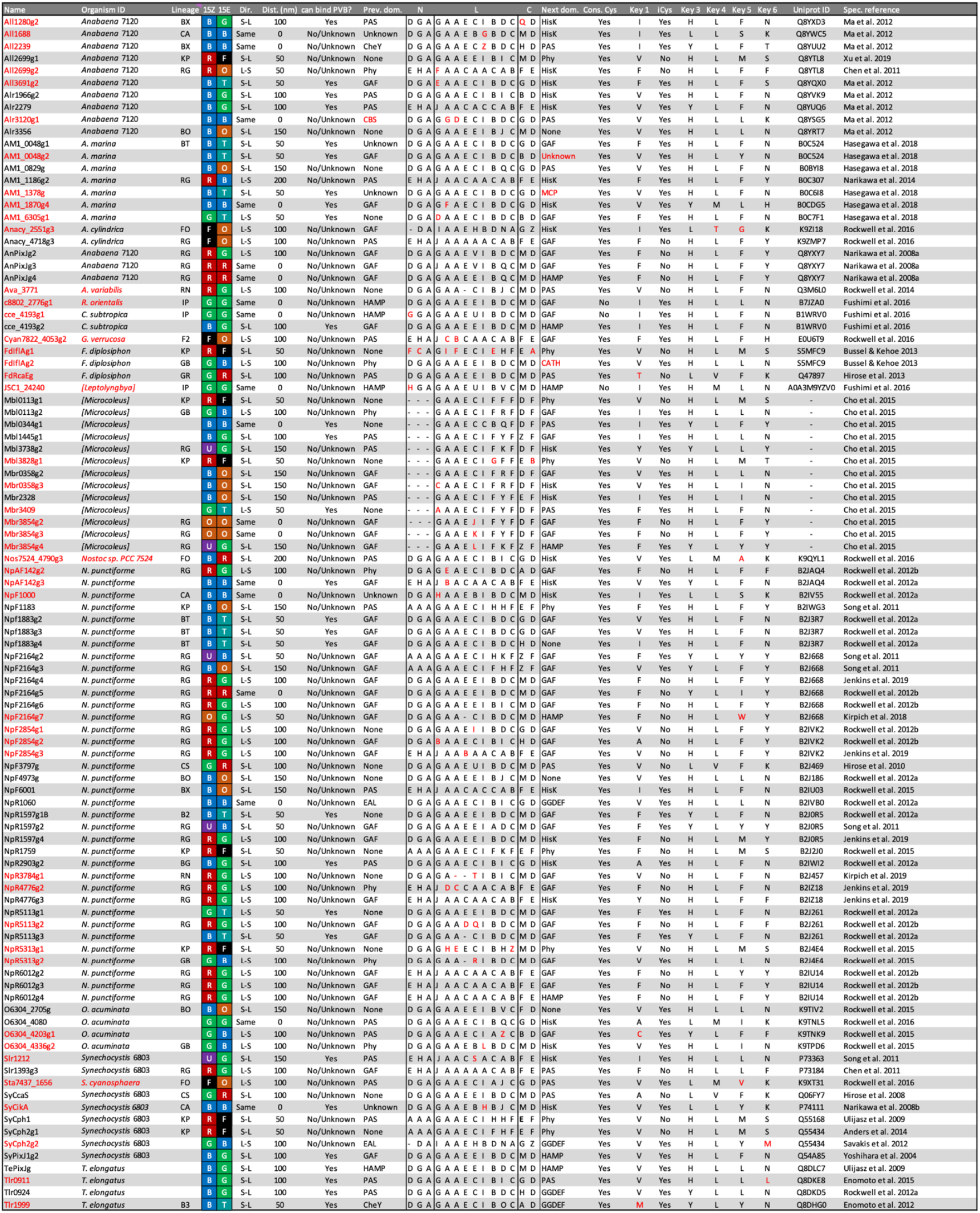
Acquired insights on all characterized GAF domains. Column 1 refers to the name of a GAF domain used throughout literature. Column 2 refers to the name of the organism encoding the GAF domain. Column 3 refers to the lineage as assigned in Figure 3.1. Columns 4,5 refers to the most readily absorbed colour in 15Z and 15E, respectively. U: UV, B: Blue, T: Teal, G: Green, O: Orange, R: Red (see Results). Column 6 refers to the change in λmax as the protein transitions from 15Z to 15E. “S-L” refers to small to large change in λmax (e.g., 400 nm -> 600 nm). “L-S” refers to large to small change in λmax (e.g., 600 nm -> 400 nm). “Same” refers to no change in λmax. Column 7 refers to the rough difference between 15Z and 15E λmax values (in nm). Column 8 summarizes known information about PVB binding (see Column 22). Column 9 shows the N-terminal neighbouring domain. The 3-letter string describing the structure of α1, α2 and α2-β1 is referred to as the “N” column. The 9-letter string describing the structure of β1-β2, β2-β3, β3, “elbow”, β3-α3, α3-β4, β4, β4-α4, α4 is referred to as the “L” column. The 2-letter string describing the structure of β6-α5 and α5 is referred to as the “C” column. Column 13 shows the C-terminal neighbouring domain. Column 14 shows whether the conserved Cys residue is present in α4 (based on ClustalW alignment). Column 15 shows Key residue 1 (based on ClustalW alignment). Column 16 shows whether a Cys residue is present in the immediate vicinity of the binding pocket (based on predicted structures). Columns 17-20 show Key residues 3-6 (based on ClustalW alignment). Column 21 lists the UniProt index of each gene. Column 22 lists the publications showing absorbance spectra and PVB-binding capacity for each gene. Characteristics unique to each category in the table are shown in red. Individual GAF domains containing unique characteristics are also shown in red.

**SUPPL. FIGURE 7:**
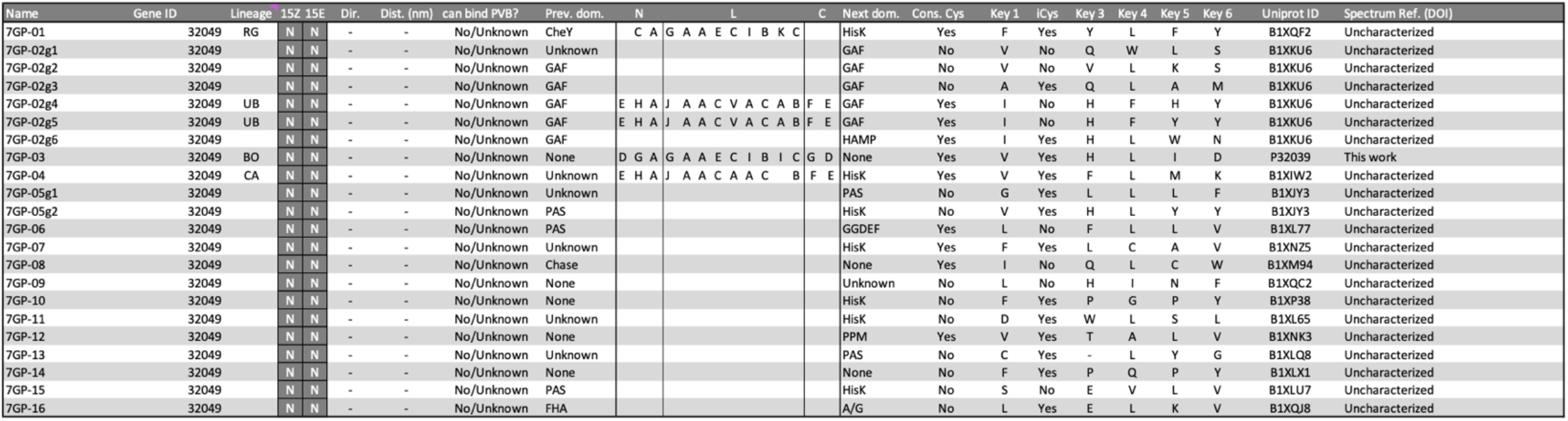
Acquired insights on all GAF domains from Synechococcus sp. PCC 7002. Column 1 refers to the name of a GAF domain used throughout literature. Column 2 refers to the Gene ID from Synechococcus sp. PCC 7002. Column 3 refers to the lineage as assigned in Figure 3.1. Columns 4,5 refers to the most readily absorbed colour in 15Z and 15E, respectively. U: UV, B: Blue, T: Teal, G: Green, O: Orange, R: Red (see Results). Column 6 refers to the change in λmax as the protein transitions from 15Z to 15E. “S-L” refers to small to large change in nm. “L-S” refers to large to small change in nm. “Same” refers to no change in nm. Column 7 refers to the rough difference between 15Z and 15E λmax values (in nm). Column 8 summarizes known information about PVB binding (see Column 22). Column 9 shows the N-terminal neighbouring domain. The 3-letter string describing the structure of α1, α2 and α2-β1 is referred to as the “N” column. The 9-letter string describing the structure of β1-β2, β2-β3, β3, “elbow”, β3-α3, α3-β4, β4, β4-α4, α4 is referred to as the “L” column. The 2-letter string describing the structure of β6-α5 and α5 is referred to as the “C” column. Column 13 shows the C-terminal neighbouring domain. Column 14 shows whether the conserved Cys residue is present in α4 (based on ClustalW alignment). Column 15 shows Key residue 1 (based on ClustalW alignment). Column 16 shows whether a Cys residue is present in the immediate vicinity of the binding pocket (based on predicted structures). Columns 17-20 show Key residues 3-6 (based on ClustalW alignment). Column 21 lists the UniProt index of each gene. Column 22 lists the publications showing absorbance spectra and PVB-binding capacity for each gene.

