## Supplementary material for "In-silico analysis of cyanobacteriochrome architectures and spectral diversity": Python script to calculate Levenstein distances

import numpy as np
def levenshtein_ratio_and_distance(s, t, ratio_calc = False):
    """ levenshtein_ratio_and_distance:
        Calculates levenshtein distance between two strings.
        If ratio_calc = True, the function computes the
        levenshtein distance ratio of similarity between two strings
        For all i and j, distance[i,j] will contain the Levenshtein
        distance between the first i characters of s and the
        first j characters of t
    """
    # Initialize matrix of zeros
    rows = len(s)+1
    cols = len(t)+1
    distance = np.zeros((rows,cols),dtype = int)

    # Populate matrix of zeros with the indeces of each character of both strings
    for i in range(1, rows):
        for k in range(1,cols):
            distance[i][0] = i
            distance[0][k] = k

    # Iterate over the matrix to compute the cost of deletions,insertions and/or substitutions    
    for col in range(1, cols):
        for row in range(1, rows):
            if s[row-1] == t[col-1]:
                cost = 0 # If the characters are the same in the two strings in a given position [i,j] then the cost is 0
            else:
                # In order to align the results with those of the Python Levenshtein package, if we choose to calculate the ratio
                # the cost of a substitution is 2. If we calculate just distance, then the cost of a substitution is 1.
                if ratio_calc == True:
                    cost = 2
                else:
                    cost = 1
            distance[row][col] = min(distance[row-1][col] + 1,      # Cost of deletions
                                 distance[row][col-1] + 1,          # Cost of insertions
                                 distance[row-1][col-1] + cost)     # Cost of substitutions
    if ratio_calc == True:
        # Computation of the Levenshtein Distance Ratio
        Ratio = ((len(s)+len(t)) - distance[row][col]) / (len(s)+len(t))
        return Ratio
    else:
        # print(distance) # Uncomment if you want to see the matrix showing how the algorithm computes the cost of deletions,
        # insertions and/or substitutions
        # This is the minimum number of edits needed to convert string a to string b
        return distance[row][col]

Morphotypes = {
"1":"AAECCBQF",
"2":"AAECIFFF",
"3":"AAECIFKF",
"4":"AAECIFRF",
"5":"AAECIFYF",
"6":"AAECIGFF",
"7":"AAEJIFYF",
"8":"AAEKIFYF",
"9":"AAELIFKF",
"10":"AAEHBDNA",
"11":"AAECIFHF",
"12":"AAECIHHF",
"13":"AAECIHKF",
"14":"A—TIBIC",
"15":"AA-CIBDC",
"16":"AA-CIBJC",
"17":"AA-RIBDC",
"18":"AADQIBDC",
"19":"AAEBGBDC",
"20":"AAEBHBJC",
"21":"AAEBIBDC",
"22":"AAEBLBDC",
"23":"AAECIAJC",
"24":"AAECIAZC",
"25":"AAECIBDC",
"26":"AAECIBIC",
"27":"AAECIBOC",
"28":"AAECIBQC",
"29":"AAEEIADC",
"30":"AAEEIBDC",
"31":"AAEEIBIC",
"32":"AAEEIBJC",
"33":"AAEEIBQC",
"34":"AAEEIBVC",
"35":"AAEIIBDC",
"36":"AAEUIBDC",
"37":"AAEVIBDC",
"38":"AAEVIBQC",
"39":"EAECIBDC",
"40":"FAECIBDC",
"41":"GDECIBDC",
"42":"HEECIBHZ",
"43":"AAAAACAB",
"44":"AABAACAB",
"45":"AACAACAB",
"46":"AACACCAB",
"47":"AACSACAB",
"48":"BACAACAB",
"49":"CBCAACAB",
"50":"DCCAACAB",
"51":"IFECIEHF",
"52":"AAEUIBVC"
}

keys_list = list(Morphotypes)

print(Morphotypes)

# main data holder
distances = []

# make rows
n = 1
while (n <= len(Morphotypes)):
	distances.append([])
	n += 1

# make columns
for each in distances:
	n = 1
	while (n <= len(Morphotypes)):
		each.append([])
		n += 1

# fill in the distances
row = 0
col = 0


for each in keys_list:
	for every in keys_list:
		distances[row][col] = levenshtein_ratio_and_distance(Morphotypes[each],Morphotypes[every])
		col += 1
	row += 1
	col = 0

# get the results

for each in distances:
	for every in each:
		print(every, end =',')
	print()

print(len(distances))

"""# calculate net divergences
net_divergence = {}

for each in range(0, len(keys_list)-1):

	counter = 0

	#horizontal sum
	for every in range(0,each):
		counter += distances[each][every]

	#vertical sum
	for every in range(each, len(distances)):
		counter += distances[every][each]

	net_divergence[keys_list[each]] = counter

print(net_divergence)"""

# testing
#print(len(Morphotypes))
#print(levenshtein_ratio_and_distance(Morphotypes["NpF1183"],Morphotypes["slr1212"]))
#print(keys_list)
#print(Morphotypes[keys_list[0]])
